# Time-restricted feeding attenuates allergic dermatitis in mice and is associated with modulation of leptin-driven inflammatory pathways

**DOI:** 10.1101/2025.10.14.680648

**Authors:** Zsófia Búr, Bernadett Vendl, Zalán Lumniczky, Bianka Farkas, Csongor G. Szántó, Domonkos Czárán, Gábor J. Tigyi, Krisztina Ella, Krisztina Káldi

**Author notes:** **Corresponding author:** Krisztina Káldi, Department of Physiology, Semmelweis University, Tűzoltó u. 37-47, Budapest, H-1094, Hungary, /ext.60411.

## Abstract

Irregular and unhealthy eating behavior contributes to the development of both metabolic and inflammatory disorders. Time-restricted eating modulates metabolism and immune function without reducing caloric intake, yet its role in inflammatory skin diseases remains unclear. Here, we examined how feeding time and diet influence contact hypersensitivity, a murine model of allergic dermatitis—a disease that affects 20% of the population. Mice were fed with normal (NC) or high-fat (HF) chow under ad libitum (AL) or 10/14 time-restricted feeding (TRF) conditions. Under AL conditions, HF diet induced metabolic dysfunction and exacerbated inflammation, characterized by increased ear swelling, leukocyte infiltration, neutrophil and cytokine accumulation, and intraepidermal pustule formation. TRF attenuated these effects and accelerated resolution, even when initiated after disease onset. HF feeding altered both the level and diurnal dynamics of serum leptin, while TRF partially restored these changes. Elevated leptin levels were associated with increased inflammatory responses including intense neutrophil accumulation and pustule formation, and both genetic and pharmacological inhibition of leptin signaling reduced disease severity. Analysis of patients’ transcriptomic data also suggested a link between leptin signaling and inflammatory skin conditions in humans. We propose that time-restricted eating beneficially influences disease progression in contact dermatitis and that leptin-associated pathways may contribute to this effect.

## INTRODUCTION

The dynamic crosstalk between metabolism and the immune system plays a crucial role in many pathological conditions. Metabolic disturbances, such as obesity, insulin resistance, and type 2 diabetes, are associated with chronic low-grade systemic inflammation. This inflammation is fueled by excess nutrient availability, which stimulates adipose tissue to produce pro-inflammatory cytokines and chemokines, promoting leukocyte recruitment and activation. Conversely, immune dysregulation can impair metabolic homeostasis and contribute to the onset and progression of autoimmune diseases, atherosclerosis, and the metabolic syndrome^1, 2, 3, 4^. These bidirectional interactions highlight the potential of dietary interventions to modulate immune responses and reduce the burden of both metabolic and immune-related diseases^5, 6, 7, 8, 9^.

Time-restricted feeding (TRF) in animal models and time-restricted eating (TRE) in humans are dietary strategies that limit food access to a specific time window of the day but do not restrict calorie intake^10, 11^. Our recent study showed that, in mice maintained on a standard diet, 10/14-hour time-restricted feeding (TRF) effectively alleviated symptoms of K/BxN serum transfer arthritis (STA)—a mouse model of rheumatoid arthritis—compared to ad libitum (AL) feeding^12^. We also found that TRF reduced the impact of a high-fat (HF) diet on STA severity^13^.

Diet-induced metabolic dysfunctions, including obesity and metabolic syndrome, are closely linked to an increased risk of inflammatory skin conditions, including contact dermatitis^14^. Allergic contact dermatitis (ACD) is a delayed-type hypersensitivity reaction of the skin, affecting around 20% of the population in Europe and North America^15, 16^. Triggered by haptens and metal ions, ACD develops in two stages^17^. The sensitization phase is clinically silent in many cases, while the elicitation phase is marked by symptoms like erythema and eczema. Contact hypersensitivity (CHS) is a well-established murine model for studying the cellular and molecular mechanisms of ACD^18, 19^. During sensitization, cutaneous dendritic cells play a key role in antigen presentation, with neutrophils and mast cells supporting T-cell priming. In the elicitation phase, keratinocytes, mast cells, and neutrophils initiate inflammation, leading to antigen-specific T and B cell activation^20, 21^. High-fat diet-induced obesity was shown to exacerbate CHS severity, but the contribution of feeding patterns to this process has not been systematically investigated ^22^.

The objective of our study was to investigate how the rhythm of feeding affects CHS under both normal and high-fat diet conditions. We observed that a relatively short period of HF diet significantly exacerbated CHS development under AL feeding conditions. Compared to AL feeding, restricting food intake to the first 10 hours of the dark phase significantly improved the inflammatory profile in both the acute and subacute forms of this disease model. We identified leptin as a candidate mediator associated with the inflammation-promoting effect of HF feeding. To explore the pathological relevance of our results, we analyzed transcriptomic data from patients with psoriasis and observed that their peripheral leukocytes expressed significantly higher levels of the leptin receptor compared with controls. Our findings and their potential translation into clinical practice may provide a basis for future investigations into therapeutic strategies for allergic dermatitis.

## METHODS

### Animals and feeding schedules

C57BL/6N (*wt*) and BKS.Cg-Dock7m^+/+^Leprdb/J male mice were kept under 12h light/12h dark cycles and fed with normal chow (NC). *Wt* male mice ranging from 60-80 days of age were also fed either high-fat (HF) diet (ssniff Spezialdiäten GmbH; NC: S8189, 17% fat; HF: D12230, 60% fat). Animals were further divided into feeding groups AL or TRF. AL groups had constant access to food, while the food access of the TRF groups was restricted to the first 10 hours of the dark (active) phase (ZT 12–22, *Zeitgeber* time, hours after light onset) (Figure 1a)^12^. Water access was unlimited and mice were housed on a grid to prevent coprophagy, with additional plastic environmental enrichment. After a 4-week dietary conditioning, experimental procedures began and the feeding protocol was maintained throughout the study.

**Figure 1.**
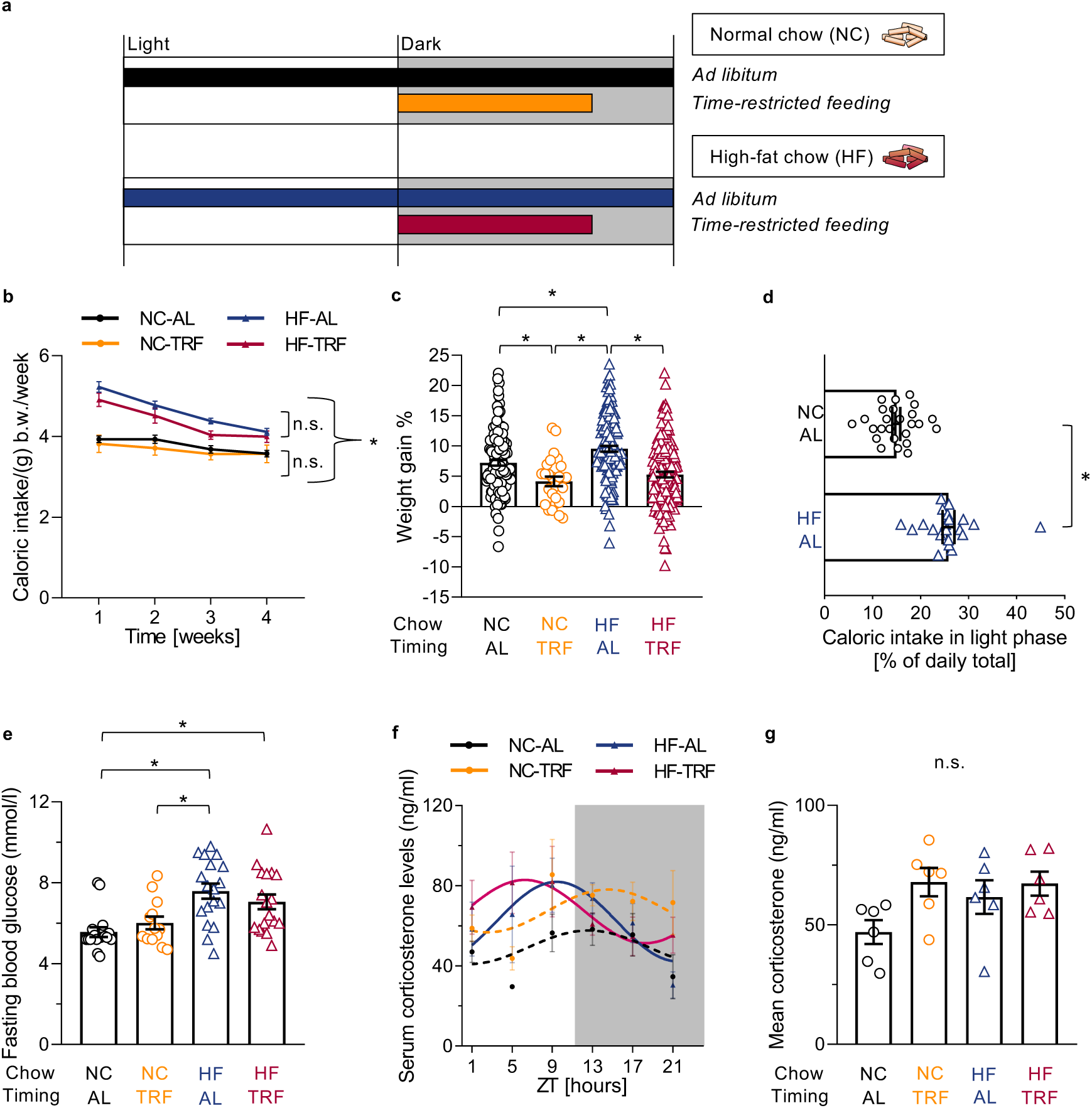
Effect of changes in feeding behavior on metabolic parameters. **a.** Schematic outline of the feeding regimes. Mice were fed either normal chow (NC) or high-fat chow (HF). *Ad libitum* (AL) fed groups had constant food availability, whereas time-restricted feeding (TRF) groups had access to food during the first 10 hours of their active phase. **b.** Caloric intake of the experimental groups during the 4-week conditioning to the feeding programs. Food consumption was recorded weekly. Data were normalized to g body weight (b.w.). Mean ± SEM, n(cages) NC-AL =23, NC-TRF=7, HF-AL=20, HF-TRF=24. Repeated Measures ANOVA, *p<0.05, significant group and group x time effect. Detailed Post Hoc results in Table S2. **c.** Body weight gain of the experimental groups after the 4-week conditioning. Data were normalized to values obtained right before the beginning of the 4-week feeding schedule. Mean ± SEM, n(NC-AL)=132, n(NC-TRF)=37, n(HF-AL)=139, n(HF-TRF)=128. One-way ANOVA, *p<0.05, significant group effect. Weekly monitored weight gain is shown in Figure S2. **d.** Distribution of caloric intake between the light and dark periods in the AL groups of animals. Percentage of caloric intake in the light period of the day was measured on each day and averaged for the 4-week period. Mean ± SEM, n(NC-AL)=25, n(HF-AL)=21, two sample t-test, *p<0.05. **e.** 4-week HF diet results in elevated fasting blood glucose levels. Mean ± SEM, n(NC-AL)=17, n(NC-TRF)=13, n(HF-AL)=18, n(HF-TRF)=18. One-way ANOVA, Post Hoc Tukey’s HSD unequal N test, *p<0.05, significant group effect. See Figure S3 for intraperitoneal glucose tolerance test. **f.** Daily changes in serum corticosterone levels in the experimental groups following the 4-week feeding period. Mean ± SEM, n/time point(NC-AL)=3-11, n/time point(NC-TRF)=3-11, n/time point(HF-AL)=3-5, n/time point(HF-TRF)=3-5. Solid lines indicate significant (p<0.05), whereas dashed lines indicate non-significant cosine curve fit. **g.** Mean corticosterone levels in the experimental groups. Daily means were calculated using data shown in Figure 1f. Mean ± SEM. One-way ANOVA, n.s not significant.

### Intraperitoneal Glucose Tolerance Test

The intraperitoneal glucose tolerance test (IPGTT) was started at ZT14 and performed as described^12^. The fasting blood glucose level corresponded to the baseline measurement point of the IGTT.

### TNCB-induced Contact Hypersensitivity (CHS)

TNCB(2,4,6-trinitrochlorobenzene)-induced CHS was provoked as described^23^. Ear thickness was measured with a caliper. Both ear thickness measurement and ear treatment were implemented under isoflurane anesthesia at ZT5. Animals were terminated by cervical dislocation at ZT5 and ears were minced for further investigations.

In the acute model, TNCB treatment of the ears was performed only once (day 6), while in the subacute model, TNCB challenges were repeated on three consecutive days (days 6, 7, and 8). To follow recovery, ear thickness was measured daily until day 20.

### Histological analysis

Ears were fixed with 4% paraformaldehyde, followed by dehydration and embedding in paraffin. 6-7 μm sections were stained with hematoxylin and eosin. Images were taken with 3DHISTECH Pannoramic^TM^ MIDI III (Pannoramic Scanner Software 5.0.0) and analyzed using SlideViewer 2.8.

### Immunohistochemistry (IHC)

To visualize neutrophils, deparaffinated sections were incubated overnight at 4 °C with anti-Gr-1 reagent diluted 1:50. The secondary antibody, Alexa Fluor™ 568 goat anti-rat IgG (Life Technologies, A11077), was diluted at 1:250 and incubated at RT for 1 h. DAPI containing mounting medium (Vector Laboratories, H-1200) was applied for nuclear staining. Images were taken using a Leica DMI6000B fluorescence microscope (Leica Microsystems).

### Analysis of leukocyte subsets

Minced ear samples were incubated in digestion solution (500 μl/sample: 200 mM HEPES (pH 7.4), 200 μg/ml Liberase™ (Roche), and 1 μg/ml DNase I (Roche) in HBSS (Gibco)) for 45 minutes at 37 °C at 1200 rpm, followed by filtering through a 70 μm strainer and rinsing with PBS. After centrifugation (5 minutes at 1000 g, 4 °C), supernatant was collected for hormone and cytokine detection, while the pellet was resuspended in PBS (5% FBS) and its cellular content was analyzed using flow cytometry (Table S1). For gating strategy, see Figure S1. Cell counts were determined using CountBright™ Plus Absolute Counting Beads (Invitrogen).

### Intradermal administration of leptin receptor blocker Allo-aca

Five days after sensitization, at ZT1, Allo-aca (10 ng/g body weight) was injected intradermally into one ear, while the other ear received vehicle injection. At ZT5, both ears were treated with TNCB. CHS severity was assessed 24 hours post-challenge.

### Measurement of hormone and cytokine levels

Leptin and corticosterone were measured in serum samples, and leptin, IL-1β, and CXCL2 levels were determined in the supernatants of ear lysates using ELISA kits (R&D Systems) according to the manufacturer’s instructions.

### Statistical analysis

Statistica software (version 14.0.1, StatSoft) was used to perform t-tests, one-way, two-way, or repeated measures ANOVA, followed by Tukey’s HSD Post Hoc test. Cosinor analysis was conducted using Matlab R2021a (MathWorks). Statistical significance was set at p < 0.05.

## RESULTS

### TRF modifies the metabolic effects of a HF diet

*wt* mice kept under 12h light/12h dark cycles were either fed *ad libitum* or subjected to a four-week feeding regimen in which food intake was limited to the first 10 hours of the dark (active) phase (TRF). The normal chow (NC) contained 17% fat and had a composition commonly used in animal facilities as a standard diet, while the high-fat chow (HF) consisted of 60 % fat (Figure 1a). Caloric intake was higher in HF than in NC groups but the difference decreased over four weeks (Figure 1b, Table S2). Importantly, the caloric intake over the 4-week period showed no significant difference between TRF- and AL-fed animals within either the HF or the NC group. However, in both diet groups, TRF animals gained less weight than AL mice (HF: p<0.001; NC: p=0.004), indicating a difference in metabolic efficiency (Figures 1c, S2, Table S3). In addition, under AL conditions, HF-fed mice consumed a higher proportion of their daily chow intake during the light phase than the NC group (p<0.001), suggesting that food composition affects the rhythm of feeding behavior (Figure 1d). To investigate metabolic regulation, intraperitoneal glucose tolerance test was performed following the 4-week feeding program. The HF diet resulted in significantly elevated fasting glucose levels compared to the NC conditions ({group effect}, p<0.001) (Figure 1e). However, the glucose tolerance test showed no difference between the groups ({group effect}, p=0.332) (Figure S3). These data indicate a partial and mild effect of HF diet on glucose homeostasis.

Corticosterone is a sensitive stress marker in mice, and its time-dependent changes provide key insights into circadian regulation affected by TRF^12^. Corticosterone concentrations remained below the stress threshold at all time points (Figure 1f, g). Moreover, high-fat conditions shifted the peak level towards the light phase of the cycle (Figure 1f, Table S4). Altogether, our data indicate that HF diet impacts both the level and the daily rhythm of glucocorticoid secretion but does not induce serious stress in the animals.

### HF diet intensifies, whereas TRF mitigates the acute CHS response

We next evaluated CHS development across experimental groups. Four weeks after initiating the diet, a subset of animals was sensitized with 3 % TNCB on the abdomen. Five days later, the ear skin was exposed to 1 % TNCB, and ear swelling was measured the next day (Figure 2a). The feeding regimen was maintained throughout the entire experiment. Sensitization resulted in a stronger swelling response to TNCB (Figure 2b) compared to non-sensitized animals (Figure S4b). HF feeding worsened, while HF-TRF mitigated swelling in both the sensitized (p<0.001) and non-sensitized groups (p=0.001) (Figures 2b and S4b).

**Figure 2.**
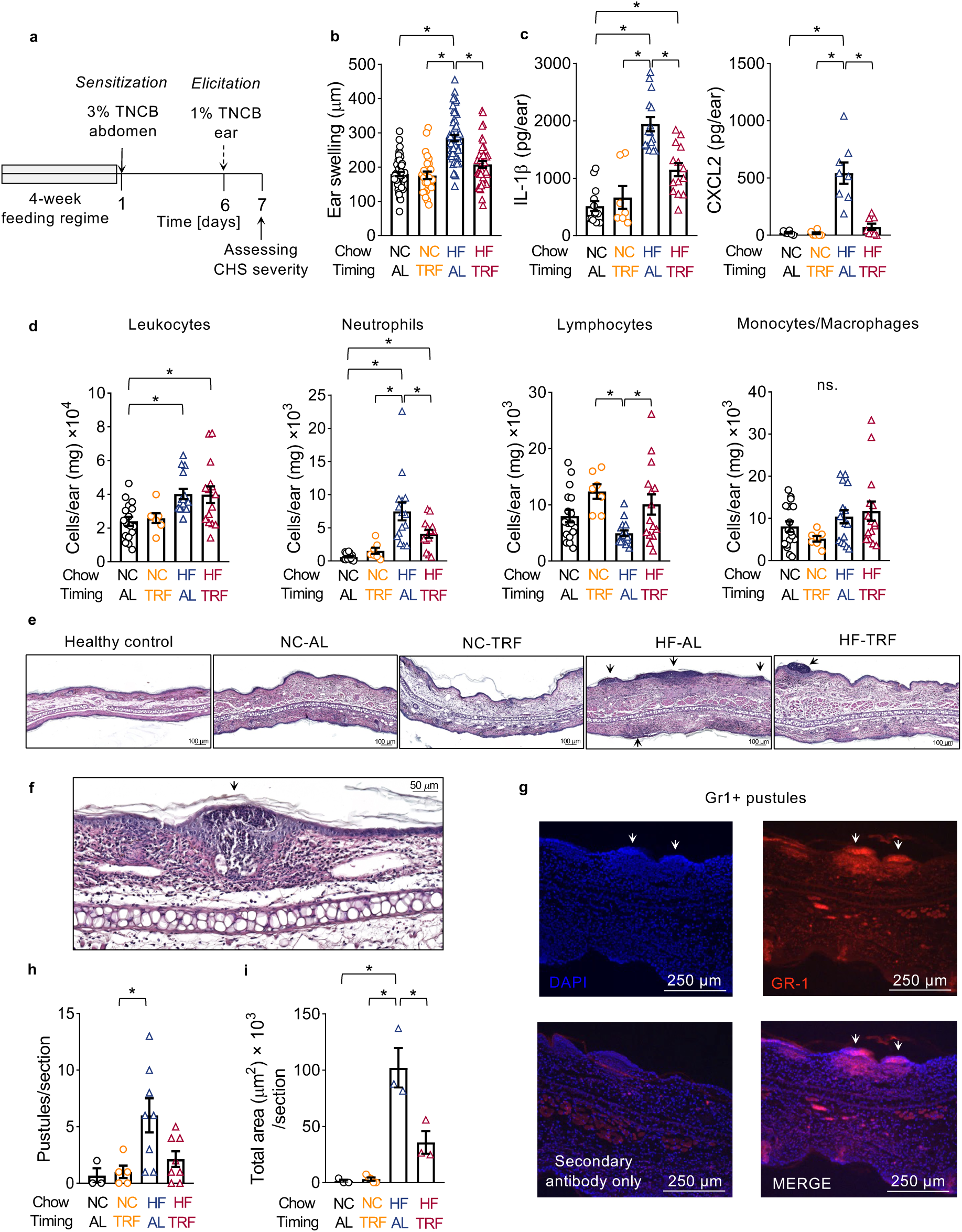
Time-restricted feeding and high-fat diet affect the symptoms’ severity of CHS. **a.** Experimental design of TNCB-induced CHS. After four weeks of conditioning in the sensitization phase, abdominal skin of mice was treated with 3 % TNCB. The elicitation phase was provoked using 1% TNCB solution applied to the ears on day 6. CHS severity was assessed 24 hours later, on day 7. **b.** Ear swelling depends on the timing and content of the diet. The ear swelling was defined as the difference in ear thickness between day 7 and day 6. Mean ± SEM, n(NC-AL)=48, n(NC-TRF)=26, n(HF-AL)=58, n(HF-TRF)=41. One-way ANOVA, Post Hoc Tukey’s HSD unequal N test, *p<0.05, significant group effect. **c.** IL-1β (left panel) and CXCL2 (right panel) levels in inflamed ears depend on diet. IL-1β: n(NC-AL)=13, n(NC-TRF)=7, n(HF-AL)=15, n(HF-TRF)=16. CXCL2: n(NC-AL)=6, n(NC-TRF)=6, n(HF-AL)=8, n(HF-TRF)=7. Mean ± SEM, One-way ANOVA, Post Hoc Tukey’s HSD unequal N test, *p<0.05, significant group effect. **d.** Accumulation of immune cells in the ears. Indicated cell counts were determined by flow cytometry. Mean ± SEM, n(NC-AL)=16-17, n(NC-TRF)=7, n(HF-AL)=15-16, n(HF-TRF)=14-15. One-way ANOVA, Post Hoc Tukey’s HSD unequal N test, *p<0.05, Leukocytes, neutrophils and lymphocytes: significant group effect, monocytes/macrophages: no significant (ns.) differences. **e.** Histological analysis of CHS development. Tissues were collected either from healthy control animals (untreated) or on day 7 (one day after TNCB treatment) from the indicated experimental groups. Hematoxylin and eosin staining of the ear section was performed. Arrows indicate pustule-like skin lesions. **f.** Intraepidermal pustule formation in HF-AL ears. Hematoxylin and eosin staining of the sections suggests neutrophil accumulation in the intraepidermal structures. **g.** Immunostaining confirms that Gr-1 (Ly6G and Ly6C) positive cells are accumulated in the pustules. DAPI staining indicates cell nuclei. **h.** HF-AL increases pustule formation. Pustules were counted on each sections. Mean ± SEM, n(NC-AL)=3, n(NC-TRF)=5, n(HF-AL)=8, n(HF-TRF)=8. One-way ANOVA, significant group effect, Post Hoc Tukey’s HSD test, *p<0.05. **i.** TRF mitigates pustule formation in HF-fed animals. Total pustular area was determined on randomly selected ear sections. Mean ± SEM, n(NC-AL)=3, n(NC-TRF)=5, n(HF-AL)=3, n(HF-TRF)=3. One-way ANOVA, significant group effect, Post Hoc Tukey’s HSD unequal N test, *p<0.05.

IL-1β and CXCL2 (MIP-2) play a central role in the activation of immune cells during the CHS response^24, 25^. Levels of both cytokines were markedly increased in the ear lysates of sensitized HF-AL mice compared to both NC groups, and the increase was significantly attenuated by HF-TRF (p<0.001, p<0.001) (Figure 2c). Consistent with changes of the cytokine levels, tissue leukocyte counts increased in sensitized HF-AL mice relative to the NC-AL group (p=0.010) (Figure 2d). Neutrophil levels changed in parallel with the total leukocyte counts, whereas lymphocytes showed opposite trends. Furthermore, TRF reduced the HF-induced elevation of neutrophil levels (p=0.030) (Figure 2d).

Histological sections of TNCB-treated ears showed signs of spongiotic dermatitis. The edema and the leukocyte infiltration of the dermis were more pronounced in the sensitized (Figure 2e) compared to the non-sensitized groups (Figure S4e). In the sensitized HF samples, the infiltration of leukocytes led to the formation of intraepidermal pustule-like lesions (Figure 2e, f). Immunostaining revealed high levels of Gr-1^+^ neutrophils in the pustules (Figures 2g). Importantly, HF-AL substantially increased pustule formation ({group effect}, p=0.013 and p<0.001) (Figure 2h, i). These structures closely resembled those observed in human pustular psoriasis and had not previously been reported in this model.

Our data suggest that even a brief, 4-week dietary intervention markedly influences CHS development. A high-fat diet worsens dermatitis, while TRF reduces disease severity.

### TRF accelerates the resolution of acute and subacute CHS in HF-fed mice and is beneficial even when started after inflammation onset

To assess whether fat content or feeding time affects the resolution of the acute skin reaction, we monitored symptoms in sensitized animals for two weeks after TNCB treatment (Figure 3a). Recovery was substantially delayed in HF-AL mice, which exhibited greater ear thickness ({group×time effect}, p=0.005) and severe macroscopic symptoms compared to the other groups (Figure 3b).

**Figure 3.**
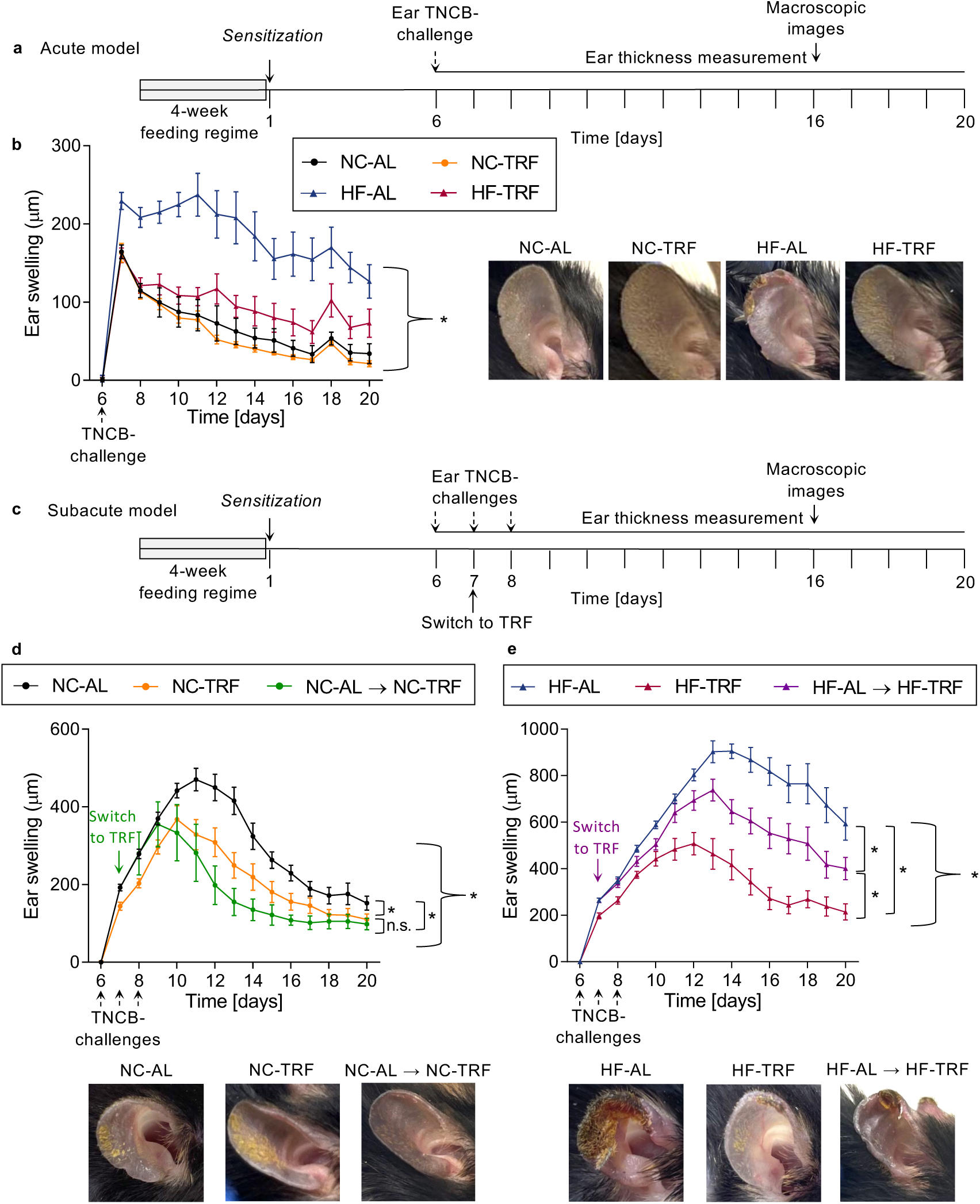
TRF exerts both preventive and therapeutic effects in acute and subacute models of CHS. **a.** Schematic outline of the experimental design in the acute model of CHS. **b.** Investigation of recovery in the acute model. Left panel: The ear swelling was defined as the difference in ear thickness between day 7 and day 6. Mean ± SEM, n(NC-AL)=8-11, n(NC-TRF)=6-9, n(HF-AL)=9-12, n(HF-TRF)=9-12. Repeated measures ANOVA (day 7 - day 20), significant group effect, *p<0.05. The data for day 7 were also used in Figure 2b. Right panel: representative macroscopic images of the ears from the indicated groups, taken 10 days after the TNCB challenge. **c.** Schematic outline of the experimental design in the subacute model of CHS. The ears were subjected to multiple TNCB treatments on the days indicated by arrows and a group of mice originally subjected to AL was switched to TRF on day 7. **d.** Both preventive and therapeutic applications of TRF improve recovery from subacute CHS in NC-fed mice. Upper panel: The ear swelling is the difference between the ear thickness on the given day and the baseline (day 6) ear thickness. Mean ± SEM, n(NC-AL)=8-17, n(NC-TRF)=7-10, n(NC-AL → NC-TRF)=3. Repeated measures ANOVA (day 8 - day 16), significant group and group x time effect, *p<0.05. The data for day 7 were also used in Figure 2b. Lower panel: representative macroscopic images of the ears from the indicated groups, taken 10 days after the TNCB challenge. **e.** HF-AL feeding profoundly delays recovery, whereas TRF – whether applied preventively or initiated after the onset of inflammation – alleviates CHS severity in the subacute model. Upper panel: The ear swelling is the difference between the ear thickness on the given day and the baseline (day 6) ear thickness. Mean ± SEM, n(HF-AL)=12-20, n(HF-TRF)=11-14, n(HF-AL → HF-TRF)=10. Repeated measures ANOVA (day 8 - day 20), significant group and group x time effect, *p<0.05. The data for day 7 were also used in Figure 2b. Lower panel: Representative macroscopic images of the ears from the indicated groups, taken 10 days after the TNCB challenge.

We next tested a subacute form of the TNCB-induced dermatitis model, similar to that used for 2,4-dinitrofluorobenzene^26^. Sensitized animals received TNCB on the ear for three consecutive days, and ear thickness was monitored over two weeks (Figure 3c). Swelling was more pronounced and prolonged than in the acute model and TRF significantly accelerated healing even with normal chow ([group effect], p=0.006) (Figure 3d). HF-AL mice had the most severe inflammation, with over fourfold ear thickening and minimal recovery by the end of the study, whereas in HF-TRF the resolution was significantly faster ([group×time effect] p<0.001) (Figure 3e). The skin’s appearance aligned with these findings (Figure 3d, e). Altogether, our data suggest that the recovery from CHS is faster in mice fed in a 10-hour period of the active phase of the day compared to AL-fed animals.

To assess whether TRF has acute effects on inflammation development and recovery, we repeated the subacute treatment with an experimental group that was fed AL before disease induction and switched to TRF following TNCB application. Under both NC and HF conditions, the AL-TRF transition accelerated the recovery, indicating that TRF exerts beneficial effects even when initiated after disease onset ([group×time effect] p<0.001; p<0.001) (Figure 3d, e). No significant difference was observed between the preventive and therapeutic application of TRF under normal chow conditions ([group effect], p=0.346). The HF-TRF and HF-AL→TRF groups differed significantly, with TRF exerting a more pronounced beneficial effect when initiated before inflammation induction ([group×time effect] p<0.001) (Figure 3d, e). Importantly, switching from AL feeding to TRF after inflammation induction attenuated the inflammation outcome under both dietary conditions.

### Leptin affects CHS severity

Beyond regulating satiety, leptin also acts as a cytokine linking metabolic disturbance to immune response^27^. Previously, we reported that TRF enhanced the rhythm and reduced the average levels of leptin RNA in visceral fat of NC animals, accompanied by corresponding changes in serum leptin levels^12^. As shown in Figure 4a, the HF diet approximately doubled the average serum leptin level under AL conditions, whereas TRF reduced this increase and shifted the peak to the dark period, matching the pattern seen with NC feeding (Figure 4a, Table S4). This indicates that TRF significantly impacts the time-dependent leptin production under HF conditions. We hypothesized that leptin as a cytokine known to affect leukocytes’ function might contribute to the exacerbation of CHS symptoms. We measured leptin levels in the inflamed ear samples and found significantly higher concentrations in HF-AL mice than in the other groups (Figures 4b, S4f). Additionally, in the HF-TRF samples, leptin levels were significantly reduced relative to the tissue lysates of the HF-AL animals in both the sensitized (p=0.039) and the non-sensitized (p=0.028) groups.

**Figure 4.**
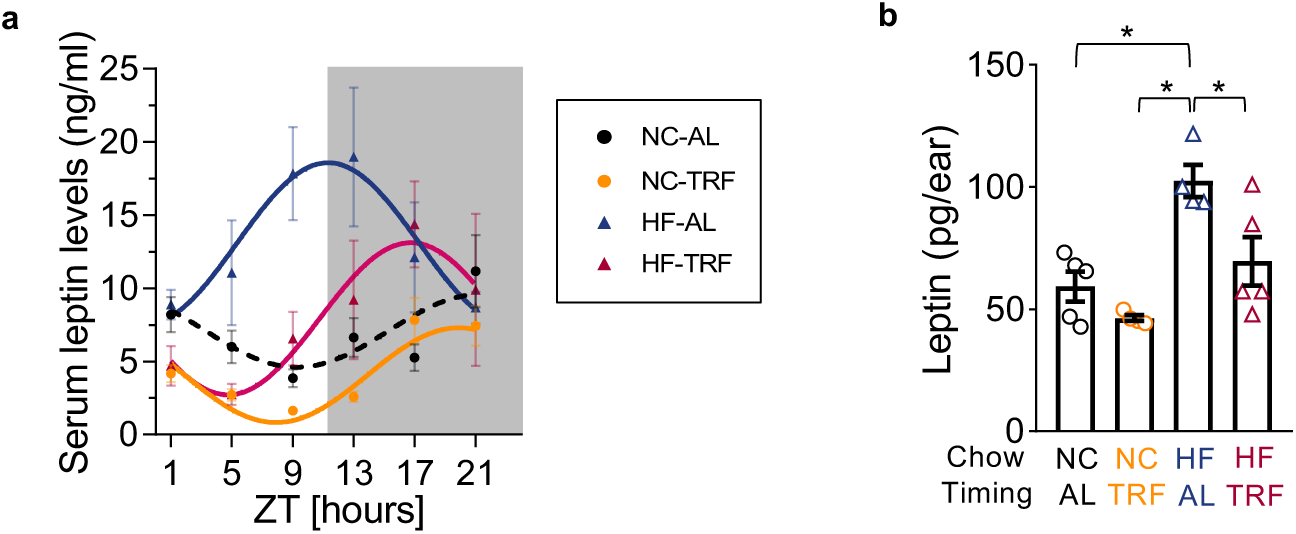
Increased leptin levels and leptin response are associated with pustule formation. **a.** Time-of-the-day-dependent changes in serum leptin levels. Mean ± SEM, n/time point(NC-AL)=3-7, n/time point(NC-TRF)=3-8, n/time point(HF-AL)=3-5, n/time point(HF-TRF)=3. Solid lines indicate significant (p<0.05), whereas dashed lines indicate non-significant (p≥0.05) fit in the cosinor model. ZT=*Zeitgeber* time (hours after the light onset). NC-AL and NC-TRF serum leptin datasets were taken from Ella et al^12^. **b.** Elevated tissue leptin levels in HF-AL mice. Tissue lysates were prepared from samples collected on day 7 of CHS and leptin levels were determined. Mean ± SEM, n(NC-AL)=5, n(NC-TRF)=4, n(HF-AL)=4, n(HF-TRF)=5. One-way ANOVA, Post Hoc Tukey’s HSD unequal N test, *p<0.05, significant group effect.

To further assess leptin’s role in the allergic response, we examined CHS development in NC-fed *db/db* mice, which lack the long form of leptin receptor (ObRb) (Figure 5a). As leptin controls appetite via ObRb, *db/db* mice are obese and produce leptin at highly elevated level^28, 29^. Immune cells involved in CHS development express either only the short form (ObRa) or both forms of the leptin receptor (reviewed in^30^), therefore, ObRa can mediate the effect of the elevated leptin levels in the *db/db* leukocytes. We found that ear swelling was more pronounced in *db/db* mice compared to *wt* controls (p=0.021) (Figure 5b), accompanied by increased production of IL-1β (p=0.057) and CXCL2 (p=0.011), prominent pustule formation (p=0.002), and enhanced neutrophil infiltration (p=0.004) (Figure 5c-e). In the next experiments, one ear of *db/db* mice was injected with the specific leptin receptor antagonist Allo-aca, while the other ear received a vehicle control. Four hours later, ears were treated with TNCB (Figure 5f). Inhibition of the leptin receptor mitigated ear swelling (p=0.024) (Figure 5g), suggesting that elevation of leptin levels could contribute to the development of severe inflammation in *db/db* mice.

**Figure 5.**
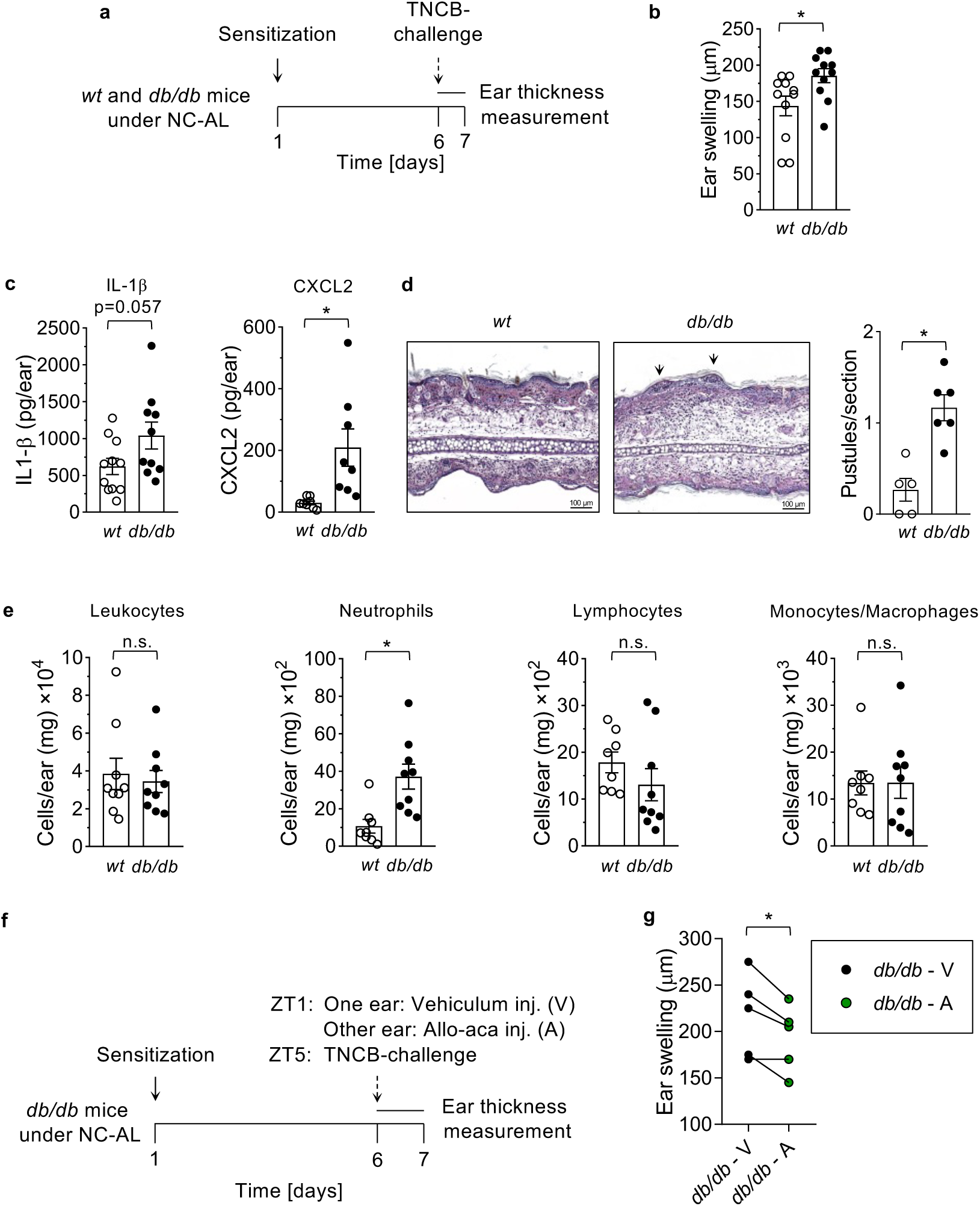
CHS severity is increased in *db/db* mice. **a.** Experimental design for induction and following CHS in the C57BLKSDock7^m^Lepr^db^ mouse strain. **b.** Relative changes of ear thickness (day 7 - day 6) of *wt* and *db/db* mice following CHS induction. Mean ± SEM, n(*wt*)=11, n(*db/db*)=12, two-sample Student’s t-test. **c.** IL-1β (left panel) and CXCL2 (right panel) levels in lysates of inflamed ears of *wt* and *db/db* mice. Mean ± SEM n(*wt*)=8-11, n(*db/db*)=8-10, two-sample Student’s t-test, *p<0.05. **d.** Increased pustule formation in *db/db* mice. Left panel: Arrows indicate pustules in the inflamed tissues of *wt* and *db/db* mice. Representative images. Right panel: Pustules were counted on sections originating from distinct samples. Mean ± SEM, n(*wt*)=5, n(*db/db*)=6 two-sample Student’s t-test, *p<0.05. **e.** Pronounced neutrophil accumulation in the ear tissue of *db/db* mice. Indicated immune cell counts were determined in lysates of inflamed ears of *wt* and *db/db* animals. Mean ± SEM, n(*wt*)=8-9, n(*db/db*)=9, two-sample Student’s t-test, *p<0.05, n.s not significant. **f.** Experimental design for the treatment with the leptin receptor antagonist. One ear of *db/db* mice was treated with vehicle, while the other ear was administered Allo-aca through intradermal injection. ZT=*Zeitgeber* time (hours after the light onset). **g.** Leptin receptor blocker, Allo-aca reduces ear swelling compared to control ears. The ear swelling was defined as the difference in ear thickness between day 7 and day 6. Data originating from vehicle- and Allo-aca treated ears of the same animal are connected. n=5, paired Student’s t-test, *p<0.05.

To examine leptin’s role in the diet-dependent inflammation control, we applied Allo-aca treatment to *wt* mice subjected to HF-AL diet (Figure 6a). Ear thickness was significantly reduced (p=0.005) (Figure 6b) and tissue levels of IL-1β and CXCL2 (Figure 6c) showed a decreasing trend in the inhibitor-treated ears relative to the vehicle-treated ones. Results of the histology also supported the anti-inflammatory effect of Allo-aca; pustules’ formation was highly inhibited by Allo-aca treatment (Figures 6d and S5). Furthermore, leptin receptor blockade decreased leukocyte infiltration (p=0.022) and subset analysis revealed that the accumulation of neutrophils (p=0.033) and monocytes (p=0.045) was significantly reduced in the inhibitor-treated ears (Figure 6e). In summary, our results suggest that leptin, as a pro-inflammatory cytokine, is associated with and may contribute to the severity of CHS.

**Figure 6.**
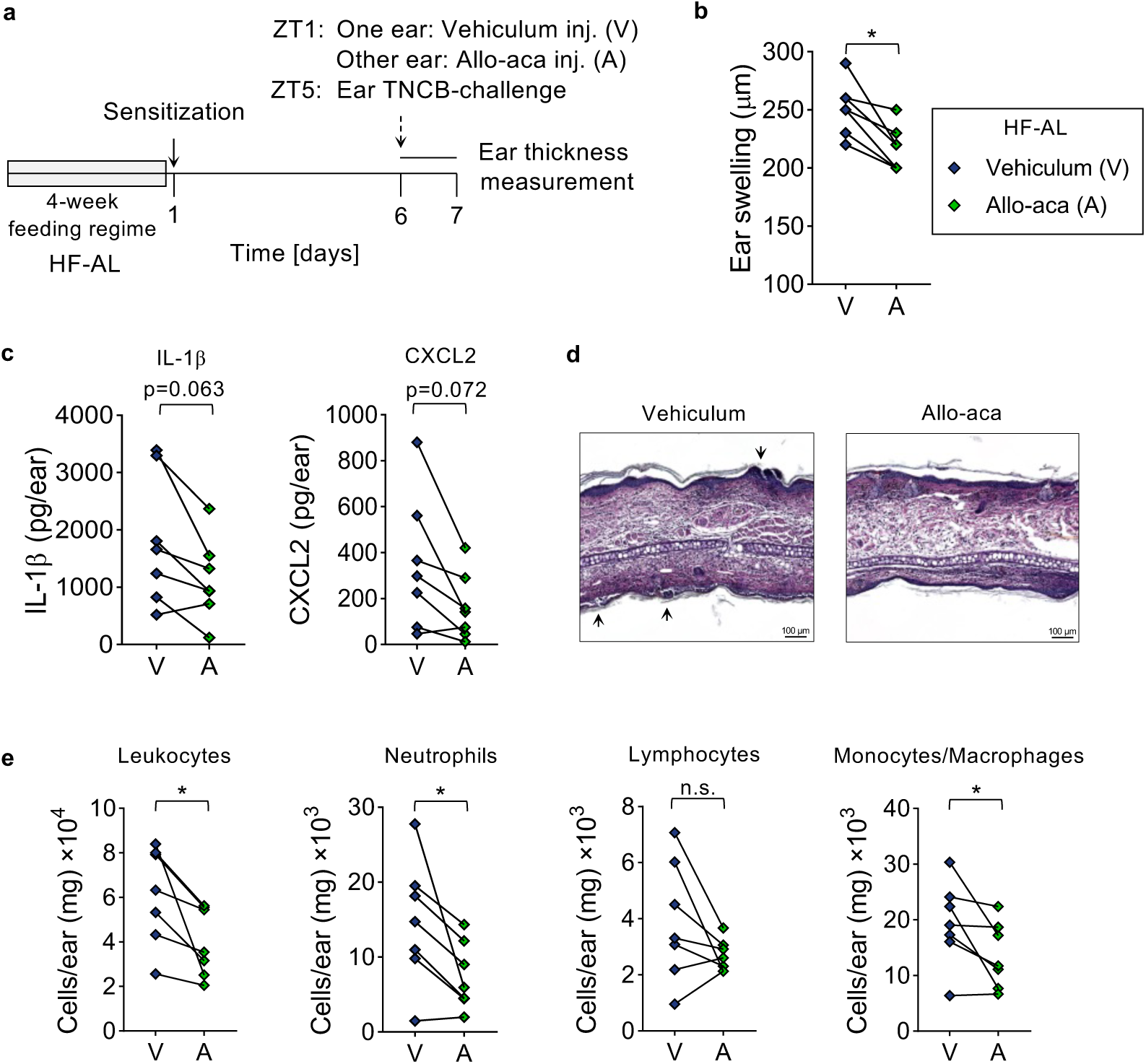
Leptin receptor blockade is associated with reduced CHS severity in HF-AL mice. **a.** Experimental design of leptin receptor blocking with Allo-aca. Allo-aca (A) or vehicle (V) were intradermally injected into the ears of HF-AL mice. ZT=*Zeitgeber* time (hours after the light onset). **b.** Ear swelling in HF-AL mice is reduced by Allo-aca. The ear swelling was defined as the difference in ear thickness between day 7 and day 6. Data originating from vehicle- and Allo-aca treated ears of the same animal are connected. n=7, paired Student’s t-test, *p<0.05. **c.** Effect of Allo-aca treatment on cytokine production in the inflamed tissues. IL-1β (left panel) and CXCL2 (right panel) levels were measured in lysates of vehicle- and Allo-aca treated ears. n=7, paired Student’s t-test. **d.** Allo-aca treatment results in decreased pustule development in CHS. Representative images of hematoxylin-eosin-stained sections of vehicle- and Allo-aca treated inflamed ears. Arrows show pustules in vehicle-treated ears. **e.** Allo-aca affects leukocyte accumulation in the inflamed ears. Counts of the indicated cell types were determined in lysates of vehicle- and Allo-aca treated ears. n=7, paired Student’s t-test, *p<0.05.

The detection of pustules resembling the histological pattern of psoriasis in the samples of HF animals was an unexpected finding. Based on the co-occurrence of pustule formation and the elevated leptin levels in our experiments, and to explore the human pathological relevance of the observations, we analyzed leptin receptor expression using transcriptomic data of PBMC samples from healthy controls and patients with generalized pustular psoriasis (GPP) or psoriasis vulgaris (GSE200977)^31^. The significantly higher transcript levels in the samples of GPP patients suggest that leptin signaling may be relevant in inflammatory skin conditions associated with pustulosis in human ({group effect}, p=0.002) (Figure 7a). Furthermore, both IL-1β receptor (*il1r1*) and *cxcr2* showed enhanced expression in the PBMC samples of patients with generalized pustular psoriasis ({group effect}, p=0.003 and 0.008, respectively) (Figure 7b, c). These findings are consistent with the elevated levels of both IL-1β and the CXCR2 ligand CXCL2 detected in the ear samples of HF-AL mice.

**Figure 7.**
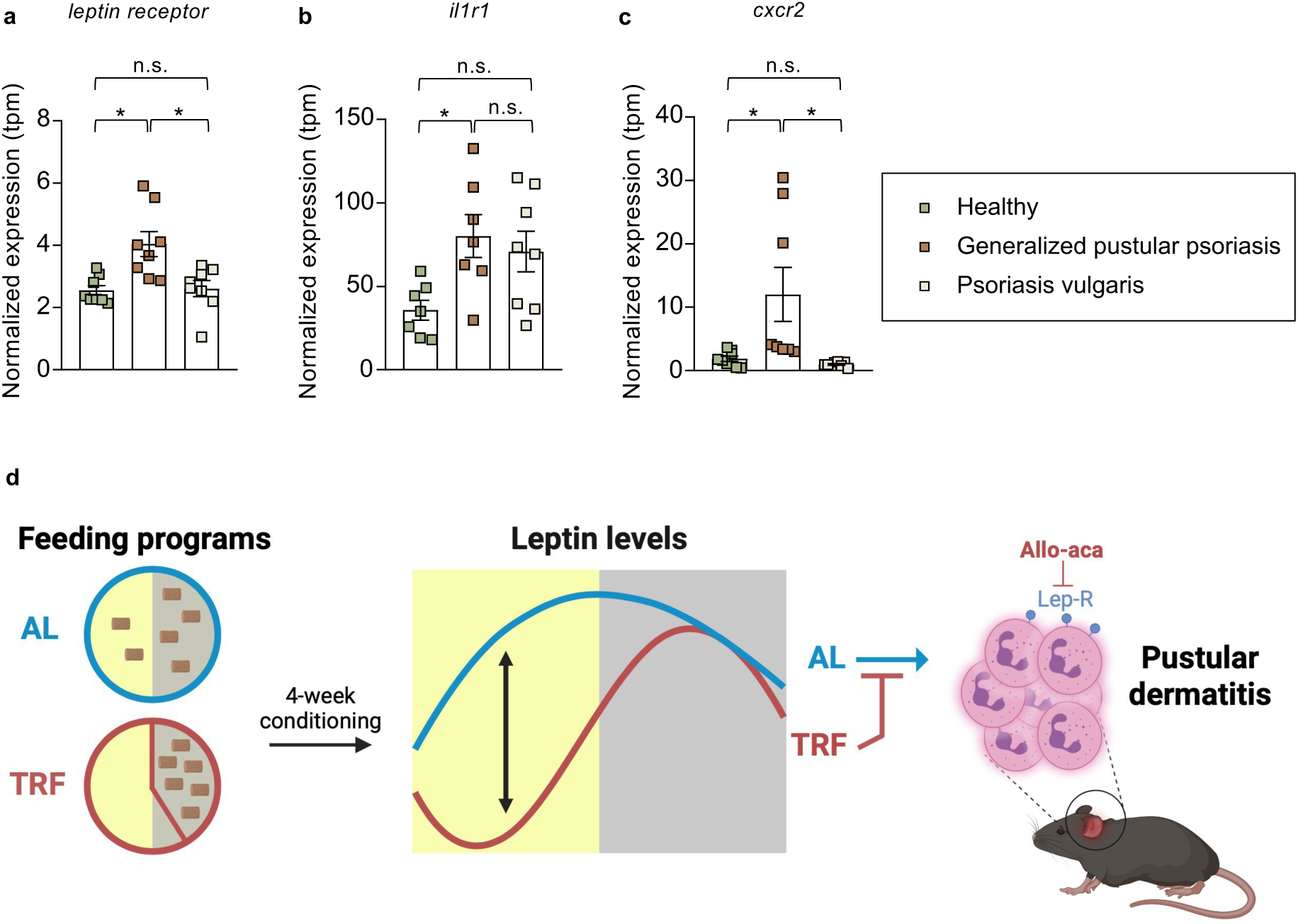
*Leptin receptor*, *il1r1* and *cxcr2* expressions are elevated in human generalized pustular psoriasis. Expression of **a.** *leptin receptor*, **b.** *il1r1* and **c.** *cxcr2* are increased in the samples of patients with generalized pustular psoriasis compared to either healthy subjects or patients with psoriasis vulgaris. Human transcriptomic data (GSE200977) were obtained from PBMC samples^31^. Expression was normalized to TPM (transcripts per million). Mean ± SEM, n=7-8, One-way ANOVA, Post Hoc Tukey’s HSD test, *p<0.05, significant group effect. **d.** Model representing the possible link between TRF and CHS outcome.

## DISCUSSION

Adverse eating behaviors - such as consuming a calorie-rich diet and maintaining irregular meal times (for example, late-evening eating) - are widespread in western societies and significantly contribute to the development of metabolic disorders, including obesity and diabetes. These pathological conditions are often associated with proinflammatory conditions leading to a higher incidence of altered immune reactions. Here we report that a relatively short period of high-fat diet elevates serum leptin levels and exaggerates pustular dermatitis; however, this worsening is markedly reduced by restricting the nutrient availability to the first 10 hours of the active period or by administering a leptin receptor blocker (Figure 7d). Detection of a large number of neutrophil-containing intraepidermal pustules in the ear sections of HF-AL animals is also a novel observation in this model. We followed the time course of symptom development for an extended period in both the acute and subacute disease models. In the subacute model, TRF promoted recovery even when mice were fed normal chow and demonstrated a pronounced effect under HF conditions, further underscoring the effectiveness of scheduled feeding in inflammation control. Moreover, TRF accelerated recovery, even when initiated after inflammation onset. However, the effectiveness of the AL-TRF switch after inflammation onset (therapeutic treatment) depended on the type of diet. While under a NC diet the preventive and the therapeutic TRF treatments were similarly beneficial, under a high-fat diet the preventive application proved to be more effective. The difference in the effectiveness of acutely applied TRF between NC and HF conditions also suggests that the composition of the diet has a significant impact on the mechanism of CHS development. A detailed understanding of the underlying pathophysiological differences between chow conditions will require further investigation. Nevertheless, our findings provide experimental evidence that TRE may influence ongoing inflammatory processes.

Unlike other studies on dietary effects on disease outcomes, our study implemented both the HF diet and TRF for only four weeks, with a relatively short and well-tolerable 14-hour daily fasting phase for TRF. As a result, the difference in the weight gain among the groups was less then 7%, suggesting that the immune effects of both the HF diet and the timely restriction of food intake are consequences of metabolic reorganization, rather than solely the chronic effects of weight change. Although considered a crude indicator, the increase in fasting blood glucose levels in the HF groups may also reflect this metabolic difference. The increased caloric intake during the light phase in HF-AL animals, compared to NC-AL mice, indicates that AL feeding more profoundly disrupts metabolic rhythm when combined with a HF diet. Interestingly, the HF diet accentuated the corticosterone rhythm, and shifted the maximum to the light phase. These results differ from a previous report^32^ which described that HF-AL diet decreased the amplitude and did not change the phase of corticosterone rhythm in the serum. As corticosterone has a permissive effect on the expression of gluconeogenetic and catabolic enzymes, advance of the corticosterone peak in the HF-TRF compared to the HF-AL group might reflect physiological adaptation of metabolic regulation to TRF.

Leptin is primarily secreted by adipocytes and circulating leptin levels directly reflect the amount of energy stored in the adipose tissue in both mice and humans^33, 34^. Leptin, through its actions on the hypothalamus, is a key regulator of appetite, however, it also affects metabolism and energy expenditure by acting on peripheral tissues, including adipose tissue, liver and pancreas^35^. As a cytokine, leptin regulates both innate and adaptive immune responses^36, 37^. Growing evidence suggests a link between increased leptin levels and an elevated risk of autoimmune diseases, such as type 1 diabetes, inflammatory bowel disease, and rheumatoid arthritis^36, 38, 39, 40^. In our experiments, the changes in serum and tissue leptin levels correlated with the degree of inflammation. Mice lacking the long form of leptin receptor exhibited enhanced symptoms of dermatitis at both the cellular and molecular levels. In the absence of ObRb, obesity develops and mice produce increased levels of leptin which might trigger accumulation and activation of neutrophils expressing the short form of the receptor (ObRa). This hypothesis was supported by the effect of the leptin receptor blocker that reduced neutrophil accumulation and cytokine levels, and alleviated both the macroscopic and microscopic symptoms of inflammation. While our data support an association between leptin signaling and CHS severity, to establish a direct causal link between leptin and the effects of TRF requires further investigations.

The histological features of the HF-AL diet-associated pustules were resembling those observed in skin samples of patients with pustular psoriasis and acute generalized exanthematous pustulosis^41^. Results of our transcriptomic analysis showing elevated leptin receptor levels in PBMCs raise the possibility that an increased leptin response of leukocytes, as well as its dietary reduction, may influence the severity of conditions associated with pustulosis in psoriasis.

The possible role of leptin signaling in increased immune responses is also supported by human data showing elevated serum leptin levels in patients with allergic contact dermatitis^42^ and in those with severe psoriasis^43^. Moreover, leptin has been shown to influence human keratinocytes and to enhance neutrophil chemotaxis^44^, which is consistent with our results.

In summary, our data suggest a model in which elevated leptin levels contribute to the severity of contact dermatitis under HF-AL conditions. TRF can mitigate both leptin production and inflammation. Moreover, using a locally applied leptin receptor blocker may represent a potential target for future therapeutic investigation.

## Ethics statement

Experimental procedures were approved by the Animal Experimentation Review Board of Semmelweis University and the Government Office for Pest County (Hungary) (Ethical approvals: PE/EA/1967-2/2017 (KA-2281) and PE/EA/00375-6/2024 (KA-4079)).

## Data availability statement

Raw data are available upon request from the corresponding author.

## Conflicts of Interest

The authors declare that the research was conducted in the absence of any commercial or financial relationships that could be construed as a potential conflict of interest.

## Author contribution

KK conceptualized the project and acquired funding. KK and KE supervised the project. KK, KE and ZB designed the experiments. ZB, BV, ZL, BF, CS, DC and KE performed the experiments. ZB analyzed the data. KK, KE, GT and ZB interpreted the results and wrote the manuscript.

## Funding information

This project was supported by the National Research, Development and Innovation Office – NKFIH projects K132393 and TKP2021-EGA-25 to KK, and by the János Bolyai Research Scholarship of the Hungarian Academy of Sciences to KE. ZB and CGS received an SE250+ Excellence PhD Scholarship (EFOP-3.6.3-VEKOP-16-2017-00009) and a scholarship of the National Academy of Scientist Education, respectively.

## Acknowledgement

We are grateful to Gábor Kovács and Éva Kemecsei for the technical support. We also thank Áron Pánczél for the contribution in implementation of intradermal injection.

## Notes

### Competing Interest Statement

The authors have declared no competing interest.

### Summary of Updates

The manuscript has been revised. Figure 7 has been expanded to include an additional panel. The supplementary materials have also been updated.

