## Supplementary material for "Time-restricted feeding attenuates allergic dermatitis in mice and is associated with modulation of leptin-driven inflammatory pathways"

**Supplementary Table 1.** List of the conjugated antibodies used in flow cytometry measurements.

**Supplementary Figure 1.** Gating strategy applied in the flow cytometry analysis of ear leukocytes.

**Supplementary Table 2.** Results of the two-way repeated measures ANOVA examining the effects of group and time on daily caloric intake.

**Supplementary Figure 2.** Body weight gain of the experimental groups during the 4-week conditioning to the feeding programs.

**Supplementary Table 3.** Results of the two-way repeated measures ANOVA examining the effects of group and time on weight gain.

**Supplementary Figure 3.** Time-restricted feeding and high-fat diet had no effect on the results of an intraperitoneal glucose tolerance test.

**Supplementary Table 4.** Cosinor analysis of serum corticosterone and leptin levels.

**Supplementary Figure 4.** Assessing CHS severity in non-sensitized mice.

**Supplementary Figure 5.** Administration of Allo-aca decreases pustule count in HF-AL mice.

**Supplementary Table 1. List of the conjugated antibodies used in flow cytometry measurements.**

| Antibody | Conjugate | Clone | Cat. No. |
| --- | --- | --- | --- |
| CD45 | PeCy7 | 30-F11 | 25-0451 |
| CD45 | PE | 30-F11 | 12-0451 |
| CD11b | eFluor450 | M1/70 | 48-0112 |
| Ly6G | FITC | 1A8-Ly6g | 11-9668 |
| Ly6G | APC | 1A8-Ly6g | 17-9668 |

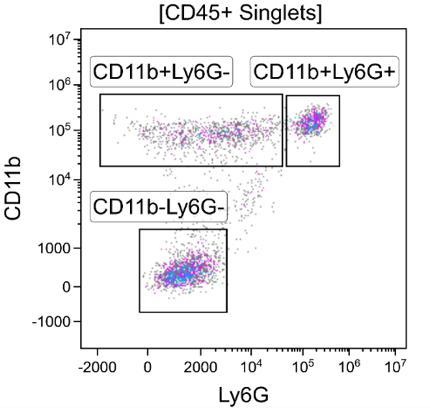

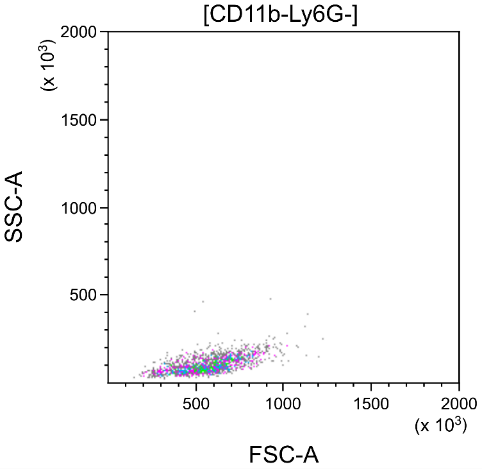

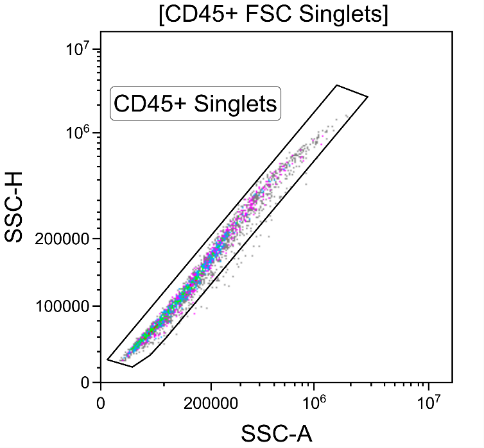

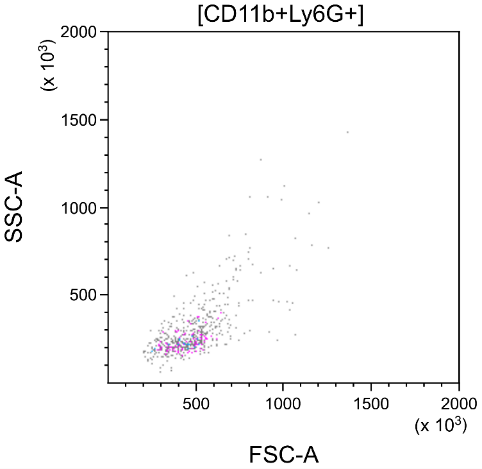

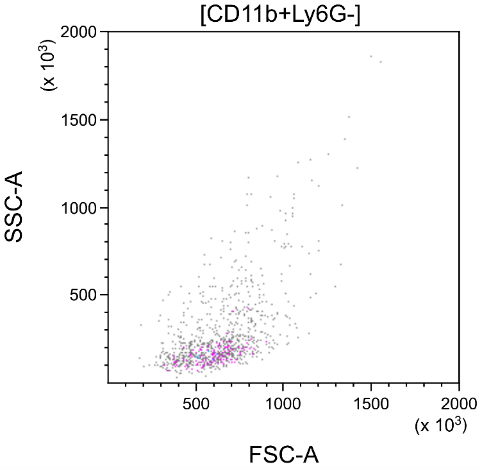

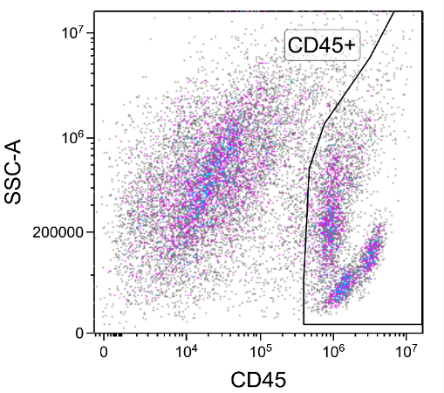

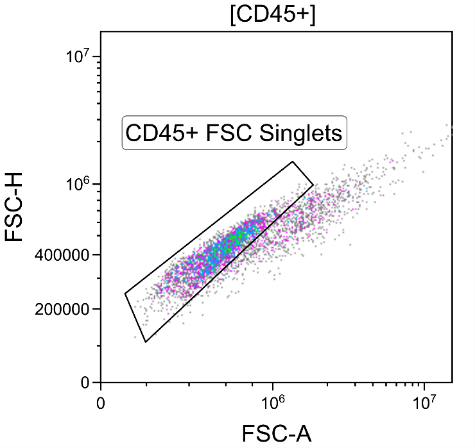

**Supplementary Figure 1. Gating strategy applied in the flow cytometry analysis of ear leukocytes.**

In CD45+ gated cells (leukocytes), singlets were gated using FSC-A and FSC-H, then SSC-A and SSC-H dot plots. In the CD45+ singlets gate, CD11b+Ly6G+ (neutrophils), CD11b+Ly6G- (monocytes and macrophages), and CD11b-Ly6G- (lymphocytes) cell populations were determined. These populations were verified using FSC-A and SSC-A dot plots. CD11b+Ly6G- gate shows heterogeneous population (different granularity and cell size) supporting that both monocytes and macrophages were included. CD11b-Ly6G- gate determines FSClow and SSClow cells, which are typical properties of small lymphocytes.

**Supplementary Table 2. Results of the two-way repeated measures ANOVA examining the effects of group and time on daily caloric intake.**

The analysis tested the main effects of group and time, as well as the group × time interaction, followed by Tukey's HSD Unequal N test. Red text indicates statistically significant difference (p < 0.05).

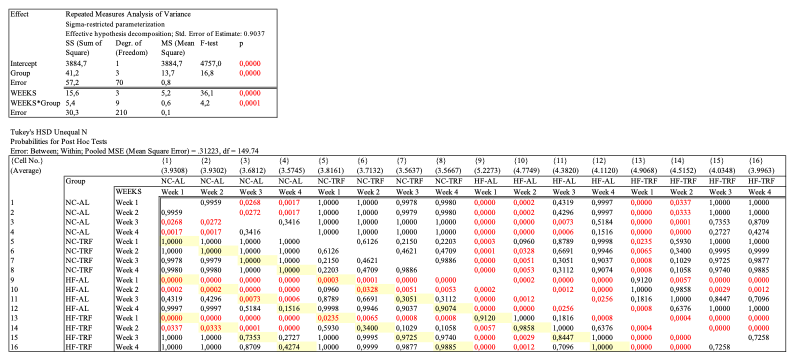

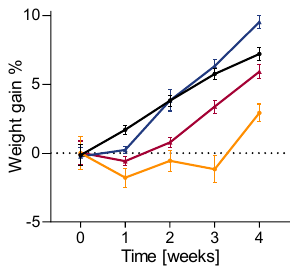

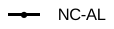

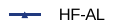

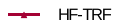

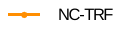

*

**Supplementary Figure 2. Body weight gain of the experimental groups during the 4-week conditioning to the feeding programs.**

Data were normalized to values obtained immediately before the beginning of the 4-week feeding program. Mean ± SEM, n(NC-AL)=131, n(NC-TRF)=37, n(HF-AL)=135, n(HF-TRF)=126. Repeated Measures ANOVA, *p<0.05, significant group and group x time effect. Detailed Post Hoc results in Table S5.

**Supplementary Table 3. Results of the two-way repeated measures ANOVA examining the effects of group and time on weight gain.**

The analysis tested the main effects of group and time, as well as the group × time interaction, followed by Tukey's HSD Unequal N test. Red text indicates statistically significant differences (p < 0.05).

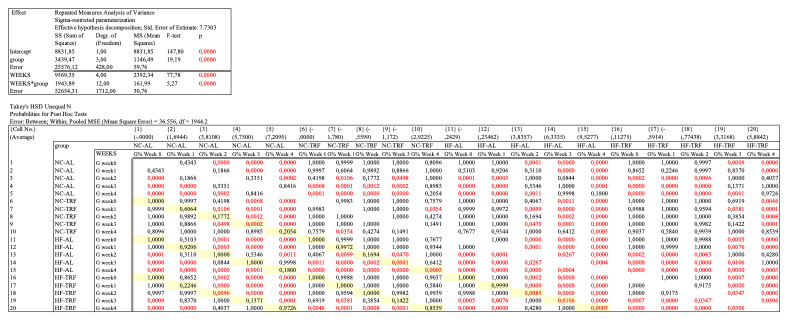

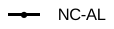

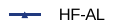

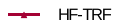

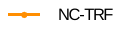

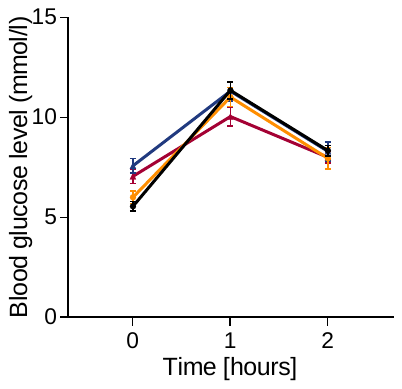

n.s.

**Supplementary Figure 3. Time-restricted feeding and high-fat diet had no effect on the results of an intraperitoneal glucose tolerance test.**

Intraperitoneal Glucose Tolerance Test (IPGTT) was performed. Mean ± SEM, n(NC-AL)=17, n(NC-TRF)=13, n(HF-AL)=18, n(HF-TRF)=18. Repeated Measures ANOVA, *p<0.05, no significant group effect.

**Supplementary Table 4. Cosinor analysis of serum corticosterone and leptin levels.**

P values in red indicate statistically significant fit (p < 0.05); p values in black indicate non-significant fit (p ≥ 0.05) in the cosinor model.

|  |  | NC-AL | NC-TRF | HF-AL | HF-TRF |
| --- | --- | --- | --- | --- | --- |
| Serum Corticosterone | Amplitude | 11.77 | 13.21 | 20.1 | 15.7 |
|  | Acrophase (ZT) | 13.38 | 14.24 | 9.6 | 6.31 |
|  | Mesor | 46.93 | 67.85 | 61.62 | 67.25 |
|  | p-value | 0.1287 | 0.182 | 0.0152 | 0.0002 |
| Serum Leptin | Amplitude | 2.66 | 3.21 | 5.59 | 5.2 |
|  | Acrophase (ZT) | 21.96 | 19.78 | 11.35 | 16.75 |
|  | Mesor | 6.85 | 4.4 | 12.92 | 7.91 |
|  | p-value | 0.0729 | 0.0039 | 0.0008 | 0.0009 |

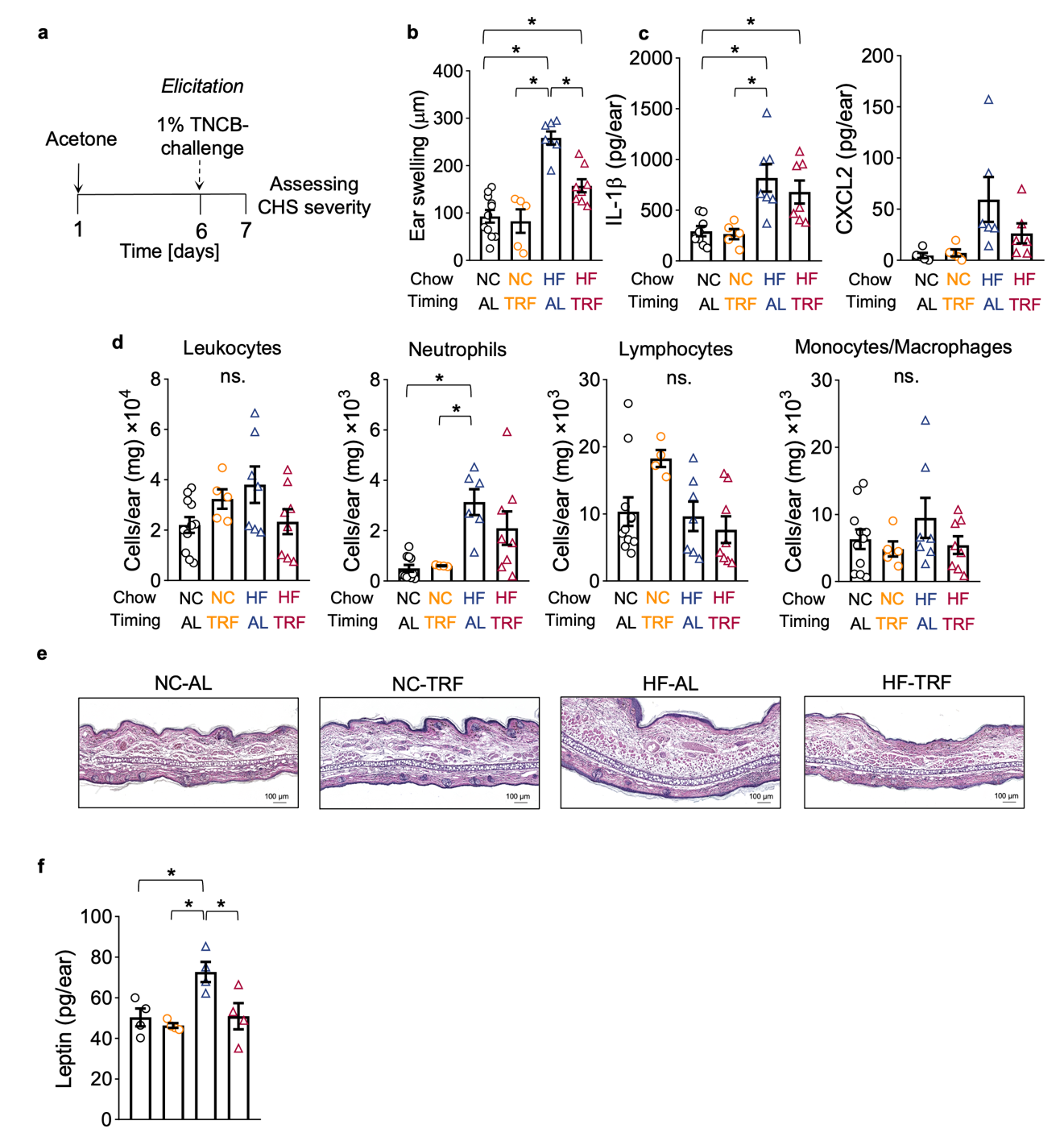

**Supplementary Figure 4. Assessing CHS severity in non-sensitized mice.**

**a.** Experimental design of TNCB-induced CHS. After four weeks of conditioning, non-sensitized animals received acetone on the shaved abdomen. Five days later, ears were challenged using 1% TNCB solution. Inflammation severity was assessed 24 hours later, on day 7.

**b.** Ear swelling depends on the timing and content of the diet. Ear thickness measured on day 7 was normalized to that measured on day 6 in the same animal. Mean ± SEM, n(NC-AL)=11, n(NC-TRF)=5, n(HF-AL)=7, n(HF-TRF)=8. One-way ANOVA, Post Hoc Tukey’s HSD unequal N test, *p<0.05, significant group effect.

**c.** IL-1β and CXCL2 levels in ear lysates are dependent on the diet. IL-1β: n(NC-AL)=8, n(NC-TRF)=5, n(HF-AL)=7, n(HF-TRF)=7. CXCL2: n(NC-AL)=5, n(NC-TRF)= 5, n(HF-AL)=6, n(HF-TRF)=6,. Mean ± SEM, One-way ANOVA, Post Hoc Tukey’s HSD unequal N test, *p<0.05, significant group effect.

**d.** Accumulation of immune cells in ear lysates of the experimental groups. Indicated cell counts were determined by flow cytometry. Mean ± SEM, n(NC-AL)=11, n(NC-TRF)=4-5, n(HF-AL)=6-7, n(HF-TRF)=8. One-way ANOVA, Post Hoc Tukey’s HSD unequal N test, *p<0.05, leukocytes, lymphocytes and monocytes/macrophages: no significant (ns.) differences, neutrophils: significant group effect.

**e.** Hematoxylin and eosin staining of the ear section. Tissues were collected on day 7 (one day after TNCB challenge).

**f.** Elevated tissue leptin levels in HF-AL mice. Tissue lysates were prepared from samples collected on day 7 of CHS. Mean ± SEM n(NC-AL)=4, n(NC-TRF)=4, n(HF-AL)=4, n(HF-TRF)=4. One-way ANOVA, Post Hoc Tukey’s HSD unequal N test, *p<0.05, significant group effect.

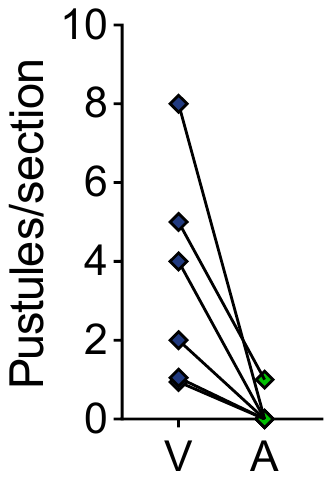

HF-AL

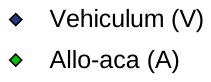

*

**Supplementary Figure 5. Administration of Allo-aca decreases pustule count in HF-AL mice.**

Data originate from two animals, and histological sections from different tissue depths were analyzed. Vehicle- and Allo-aca treated ears of the same section are connected. n=6, paired Student’s t-test, *p<0.05.
